# Global distribution and diversity of alien Ponto-Caspian amphipods

**DOI:** 10.1101/2021.07.19.452907

**Authors:** Denis Copilaș-Ciocianu, Dmitry Sidorov, Eglė Šidagytė-Copilas

## Abstract

The Ponto-Caspian region is an important donor of aquatic alien species throughout the Northern Hemisphere, many of which are amphipod crustaceans. Despite decades of ongoing spread and negative effects on native biota, a complete picture of the global diversity and distribution of these amphipods has yet to emerge, hampering efficient monitoring and predictions of future invasions. Herein, we provide a comprehensive summary of alien species taxonomic and ecomorphological diversity, as well as high-resolution distribution maps and biogeographical patterns based on >8000 global records. We find that up to 39 species in 19 genera and five families, belonging to all four currently recognized ecomorphs, are potentially alien, their diversity gradually decreasing with distance from the native region. Most species (62%) have limited distributions, 15% are widespread, and 23% exhibit intermediate ranges. We also find that regions adjacent to the native areal are comparatively less well-sampled than more distant regions. Biogeographical clustering revealed three faunal provinces that largely correspond with the Southern, Central and Northern invasion corridors. We conclude that 1) alien amphipods are a representative subsample of the native Ponto-Caspian phylogenetic and ecomorphological diversity, and 2) that their biogeographical patterns are driven by anthropogenic factors acting on distinct native regional species pools.

## Introduction

The Ponto-Caspian region harbors a unique fauna adapted to withstand a broad regime of salinity conditions (Reid and Orlova 2002). Human mediated dispersal via shipping, artificial canals, intentional introductions, and increased ionic content in inland water has enabled some of these species to disperse throughout the Northern Hemisphere as far west as the North American Great Lakes, and east into Central Asia, sometimes with enormous economic and ecological consequences (Vanderploeg et al. 2002; Strayer 2009). Among these invasive Ponto-Caspian species, amphipod crustaceans are the most numerous and diverse (Fig. 1) (Bij de Vaate et al. 2002; Cuthbert et al. 2020).

**Fig. 1.**
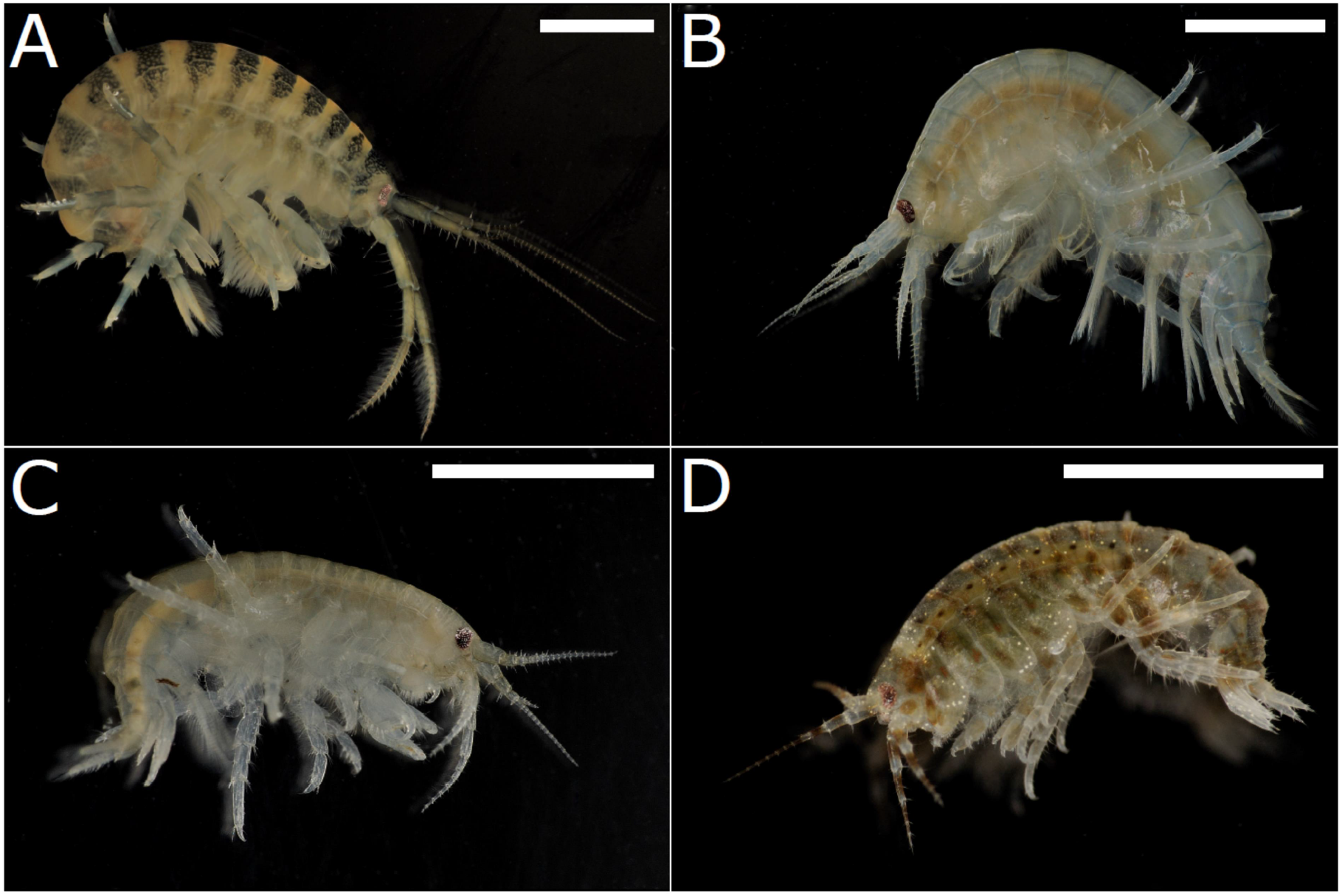
Ecomorphological diversity of Ponto-Caspian gammarid amphipods. A) *Dikerogammarus villosus* (crawler ecomorph), B) *Pontogammarus robustoides* (digger ecomorph), C) *Obesogammarus crassus* (digger ecomorph), *D) Chaetogammarus warpachowskyi* (clinger ecomorph). Scale bar = 0.5 cm. Photographs Denis Copilaș-Ciocianu.

Invasions of alien Ponto-Caspian amphipods (hereafter APCAs) are often followed by extinction of native species, and restructuring of ecological communities (Dermott et al. 1998; Van Riel et al. 2006; Arbačiauskas 2008; Grabowski et al. 2009; Arbačiauskas et al. 2017). It is generally thought that native species replacement is due to the higher aggressiveness, reproductive output and adaptability of the Ponto-Caspian invaders (Dick and Platvoet 2000; Grabowski et al. 2007; Arbačiauskas et al. 2013; Bacela-Spychalska and Van Der Velde 2013; Šidagytė and Arbačiauskas 2016). However, relatively few alien Ponto-Caspian amphipods have been studied thus far, most attention being focused on a handful of the most widespread species.

The dispersal of APCAs has a long history dating back more than a century. Their range expansions have been primarily facilitated by the opening of artificial canals in the 18^th^ century, which connected the river basins of Central/Western Europe to those of the Black Sea (Bij de Vaate et al. 2002). Some of the earliest species known to have expanded their ranges are *Chaetogammarus ischnus* and *Chelicorophium curvispinum*, possibly as early as the late 19^th^ century (Jarocki and Demianowicz 1931; Crawford 1935). A second important wave of invasions took place in the 1960s where a number of species were deliberately introduced and successfully acclimatized in reservoirs and lakes in the Baltic region, from where they subsequently expanded (Bij de Vaate et al. 2002; Arbačiauskas et al. 2017). At the same time, similar patterns of deliberate introductions took place also in other areas of the former Soviet Union, especially in newly built reservoirs on the Dnieper, Volga and Kura rivers (Ioffe 1973; Jazdzewski 1980; Grigorovich et al. 2002). The opening of the Rhine-Main-Danube canal in 1992 has promoted the dispersal of many amphipods into Western Europe (Bij de Vaate et al. 2002). From here, *C. ischnus* has even been transferred into the Great Lakes region in North America through shipping activity, where it successfully established along with other Ponto-Caspian species (Witt et al. 1997; Cristescu et al. 2004). Several species are already established in the British Isles as well (MacNeil et al. 2010). Nowadays, the distribution of APCAs in Eurasia stretches from Ireland to eastern Kazakhstan, where they are common in most of the large rivers and their estuaries.

Despite their wide distribution and documented negative effects, few studies have previously documented the distribution of APCAs at the continental scale (Jazdzewski 1980; Barnard and Barnard 1983). However, these studies are decades old, and, since then, the ranges of these species have increased dramatically. Thus, to the extent of our knowledge, there are no high resolution, up to date, distribution maps comprising the global ranges of APCAs, despite that numerous studies have provided such data at a regional scale (Jażdżewski and Konopacka 1988; Bollache et al. 2004; Grudule et al. 2007; Žganec et al. 2009; Arbačiauskas et al. 2011; Borza 2011; Borza et al. 2015; Meßner and Zettler 2018), and range expansions are routinely reported (Copilaș-Ciocianu and Arbačiauskas 2018; Moedt and Van Haaren 2018; Minchin et al. 2019; Lipinskaya et al. 2021). Even though the spatio-temporal dispersal patterns are relatively well known for many species (Bij de Vaate et al. 2002; Grigorovich et al. 2002; Cristescu et al. 2004; Rewicz et al. 2015; Jażdżewska et al. 2020), a lack of high resolution distribution data precludes efficient global monitoring and hampers predictions of future invasions. Such predictions are crucial for an effective management aimed at preventing further dispersal (Gallardo and Aldridge 2015).

Apart from sparse knowledge on distribution ranges, it is also unclear how many APCAs occur outside their native area. To the extent of our knowledge, a comprehensive list of these species along with their taxonomic and ecomorphological diversity, as well as modes of introduction has never been compiled. Such information is, again, of high relevance in forecasting future invasions (Borza et al. 2017).

Considering the above shortcomings, with this paper we aim to compile for the first time information regarding the diversity and global distribution of all Ponto-Caspian amphipods that are known to occur outside their native range. Specifically, we aim to bring insight into patterns of taxonomic and ecomorphological diversity, as well as a better understanding of biogeographical patterns based on up-to-date, high resolution distribution data.

## Material and Methods

### Defining the native range

The boundary between the native and non-native areas of a species’ distribution is often not straightforward to define (Courchamp et al. 2020; Pereyra 2020). We acknowledge that a single definition applicable to all APCA species is difficult given the high taxonomic and ecological diversity of the group (Copilaș-Ciocianu and Sidorov 2021). Many species have well-documented introductions (Grigorovich et al. 2002, 2003; Arbačiauskas et al. 2011) and spatio-temporal dispersal patterns (Bij de Vaate et al. 2002) outside the native area, while in some cases it is assumed that they dispersed since the Late Pleistocene (Cristescu et al. 2004). As such, we aim for an operational definition that is generally applicable.

We consider that the native range of APCAs encompasses the seas themselves (Black, Azov, Caspian and Aral) as well as the adjacent lagoons, river mouths and deltaic/estuarine regions. This is mainly because Ponto-Caspian amphipods have inhabited and diversified in this area (i.e. Paratethys Sea) since the late Miocene, as evidenced by fossils as well as time-calibrated phylogenies (Derzhavin 1927; Hou et al. 2014; Hou and Sket 2016; Copilaș-Ciocianu et al. 2020; Copilaș-Ciocianu and Sidorov 2021). Only four non-alien species that belong to the Ponto-Caspian amphipod group (ca. 4%) are found exclusively outside the Ponto-Caspian realm, all of these being found in the nearby Balkan Peninsula or the Caucasus (Copilaș-Ciocianu and Sidorov 2021). Therefore, our definition is conservative as it does not include lower sections of rivers that stretch further than the deltaic regions (Gogaladze et al. 2020). Drawing a line between the native and invasive areal is not straightforward in such linear habitats (Pereyra 2020). Likewise, setting a fixed threshold distance is not feasible as each river has its idiosyncrasies.

### Taxonomic framework

As a taxonomic framework we used a recent review of Ponto-Caspian amphipod diversity (Copilaș-Ciocianu and Sidorov 2021). Two species were found to be taxonomically ambiguous and should be treated with caution. First, we do not consider that the north-western Black Sea (including Central and Western Europe) populations of *Trichogammarus trichiatus* are conspecific with the eastern Black Sea ones (Caucasus region) based on which this species was originally described (Martynov 1932). The north-western populations were initially described as *Chaetogammarus tennelus major* by Caraușu (1943) from lagoons in Romania and Bulgaria, and were later synonymized with *T. trichiatus* (Dedju 1967; Straškraba 1969), a situation that is currently accepted (Rachalewski et al. 2013). However, unpublished molecular and morphological data confirms the distinctiveness of both taxa, which are not even closely related (Copilaș-Ciocianu in prep.). As such, all the non-Caucasian populations are provisionally referred to as *Trichogammarus* cf. *trichiatus*, pending further study. Second, we do not distinguish between the records of *Yogmelina pusilla* and *Y. limana*, since the latter has never been reported outside its type locality, probably due to misidentification with the former (Karaman and Barnard 1979).

### Data collection

We collected data from any available source (publication, conference proceedings, report or online database) which provided precise locality information (either coordinates or locality name). Our own unpublished data was included as well. Whenever possible, data was also extracted from maps (∼10 km accuracy) if locality information was not provided in the text. We generally focused on collecting data from non-native areas. However, for the most widespread species we also collected all available data from their native ranges as well.

Data from online databases was obtained from GBIF (https://www.gbif.org/; https://doi.org/10.15468/dl.iddp60; https://doi.org/10.15468/dl.uvqzia; https://doi.org/10.15468/dl.ythkoe; https://doi.org/10.15468/dl.qrkgyj; https://doi.org/10.15468/dl.i8uafv; https://doi.org/10.15468/dl.h40xk1; https://doi.org/10.15468/dl.agjzc4; https://doi.org/10.15468/dl.kkgszv; https://doi.org/10.15468/dl.xmtnxw) and the Czech Hydrometeorological Institute (http://hydro.chmi.cz/). Data collection ended in March 2021.

On each species we collected further data regarding its mode(s) of introduction, ecomorphology and taxonomy. Following Grigorovich et al. (2002) we distinguished five types of species introduction modes: deliberate, accidental, facilitated by hydrotechnical constructions (canals and reservoirs), shipping, and natural dispersal. Ecomorph and taxonomic assignment followed Copilaș-Ciocianu and Sidorov (2021). The corophiid genus *Chelicorophium* was not included in the ecomorphological classification as ecomorphs have been established only for gammaroidean Ponto-Caspian amphipods (Copilaș-Ciocianu and Sidorov 2021).

### Data analysis

We examined the relationship between the total number of species and total number of alien species within a genus using Pearson correlation. We also tested whether the number of species within ecomorphs statistically differs between the native and non-native regions. For this we used Fisher’s exact test with 9999 permutations. Statistical analyses were performed with PAST 3 (Hammer et al. 2001).

We explored to what extent the phylogenetic diversity of the alien species represents the total endemic diversity of gammaroidean and corophioidean Ponto-Caspian amphipods. For this we used the PhyRe python script (Plazzi et al. 2010), running the analysis at the species level with 1000 random replicates and default settings. For each species we also added the superfamily, family and genus level. The master list with the reference taxonomy followed Copilaș-Ciocianu and Sidorov (2021).

Distribution maps were created with QGIS 3.10 (QGIS Development Team 2016). We estimated the number of alien species per country and per freshwater ecoregion (Abell et al. 2008) with the help of layers downloaded from https://geospatial.tnc.org/. Species richness and sampling redundancy grids were created with Biodiverse 2.0 software (Laffan et al. 2010) using 1° cells. Biogeographical regionalization was performed using the Infomap Bioregions server (https://www.mapequation.org/bioregions/) (Edler et al. 2017). The analysis was run with default settings, except the number of trials which was set to 10 (maximum) and the value of the cluster cost parameter was set to 1.2. This was the most conservative value as increasing it resulted in only one biogeographical region. For the last two analyses only records from the non-native regions were used.

## Results

Overall, we reviewed 527 sources, of which 322 had precise locality information (supplementary references will be provided during the review process). Overall, we obtained 8145 distribution records. Data collection ended in March 2021. The full dataset is available at Figshare (doi will be provided after acceptance).

### Biodiversity and modes of introduction

In total, we consider that up to 39 species (in 19 genera and five families) are potentially alien, as they were reported occurring outside their native region (Table 1). This number represents approximately 40% of all currently known Ponto-Caspian endemic amphipod diversity (96 species, Copilaș-Ciocianu and Sidorov 2021).

**Table 1.**
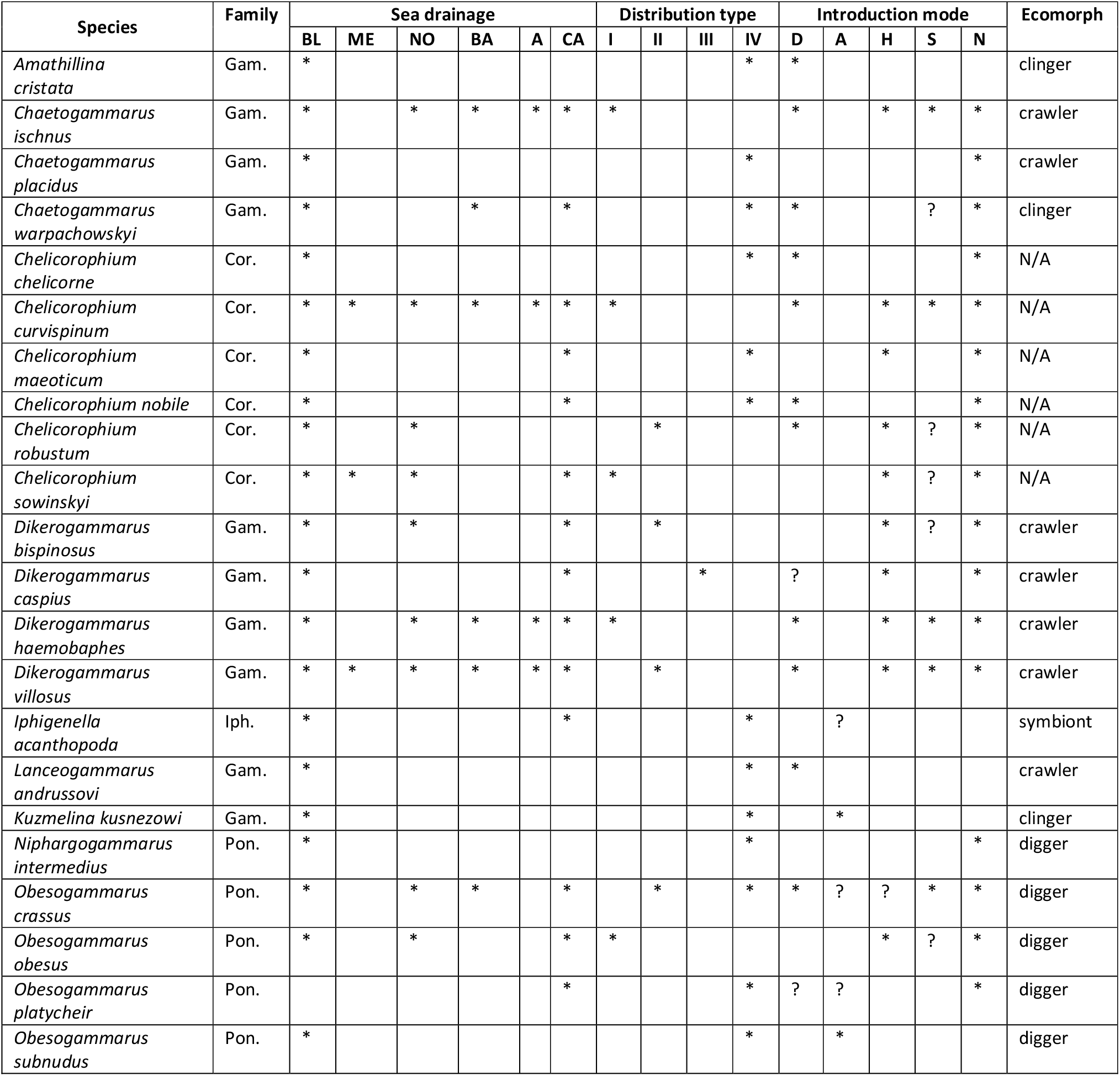
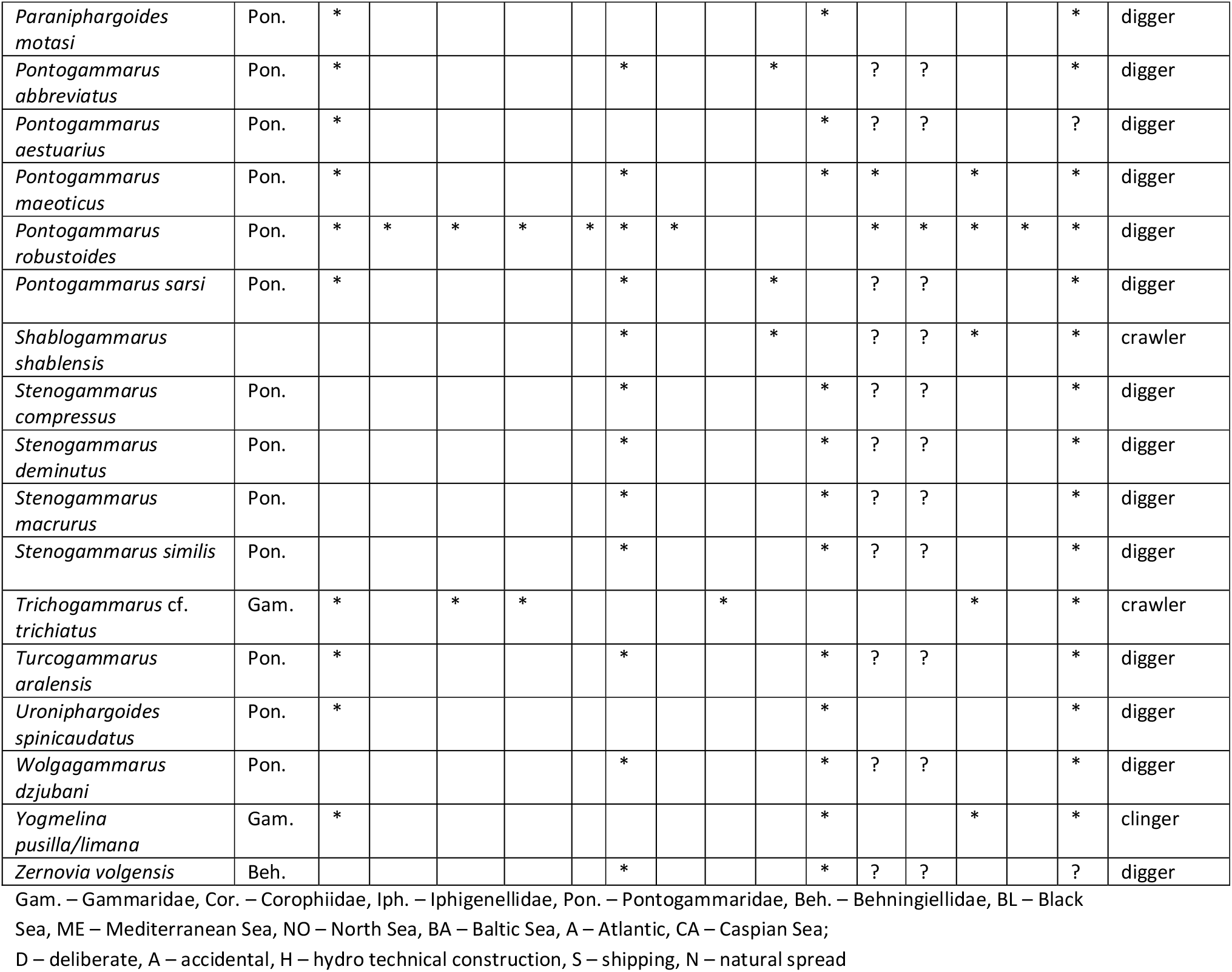
Checklist of alien Ponto-Caspian amphipods, their non-native distribution, biogeographical pattern, mode of introduction and ecomorphological classification.

We identified a strongly positive correlation between the total number of species and alien species within a genus (Pearson correlation, r^2^ = 0.81, p = 0.0001) (Fig. 2A). With respect to ecomorphology, we find that 19 species are diggers, 9 are crawlers, 4 are clingers and one is a symbiont (Fig. 2B, Table 1). We did not find significant differences between the number of non-native and native species within ecomorphs (X^2^ = 1.148, p = 0.76), meaning that the ecomorphological diversity in the non-native area is similar to that in the native area.

**Fig. 2.**
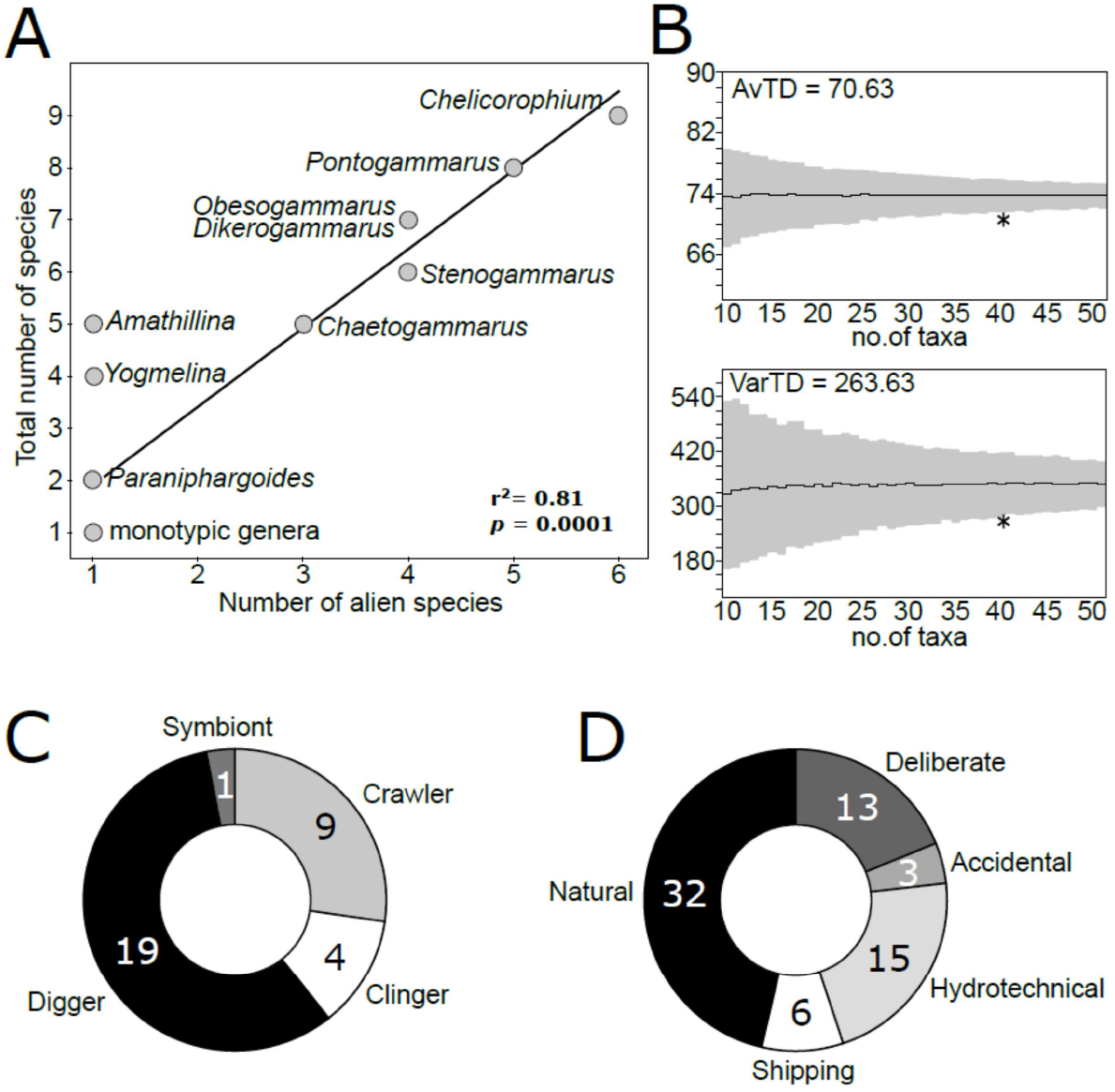
A) Relationship between the total number of species and number of invasive species within a genus. B) Funnel plots indicating the average taxonomic distinctness (AvTD) and variation in taxonomic distinctness (VarTD) of alien Ponto-Caspian amphipods. Asterisks indicate the empirical parameter values, thick black lines indicate mean, and gray area reflects the 95% confidence interval of the 1000 replicates. C) Pie-chart indicating the number of species that belong to each of the four currently recognized ecomorphs. D) Pie-chart indicating the number of species that have expanded their ranges via different introduction modes. Note that one species may have dispersed via several modes.

The phylogenetic representativeness analysis revealed that the alien species represent a relatively good proportion of the total phylogenetic diversity of the endemic gammaroidean and corophioidean amphipod fauna. The average taxonomic distinctness (AvTD = 70.63) and variation in taxonomic distinctness (VarTD = 263.63) were slightly below the lower bound of the 95% interval, the former indicating modest representativeness while the latter indicating good representativeness (Fig. 2B). Moreover, von Euler’s index of imbalance (= 0.19) was well below the recommended threshold of 0.25, which also indicates good representativeness (Plazzi et al. 2010).

Regarding introduction mode, we find that up to 32 species seem to have dispersed naturally (Fig. 2C). However, this dispersal type is almost always associated with human activity (Table 1). A total of 16 species have been documented as introduced (13 deliberately and 3 accidentally), 15 have spread via hydrotechnical constructions (reservoirs and canals), and 6 have spread through shipping activity (Fig. 2C). For a number of species, the exact dispersal mode is still uncertain (Table 1).

### Biogeography

Distribution maps of all 39 species are presented in Figures 3-5. For the 14 most widespread species we present records spanning their entire distribution area, including the native region (Figs. 3-4). For the reminder of species we present only non-native distribution records (Fig. 5).

**Fig. 3.**
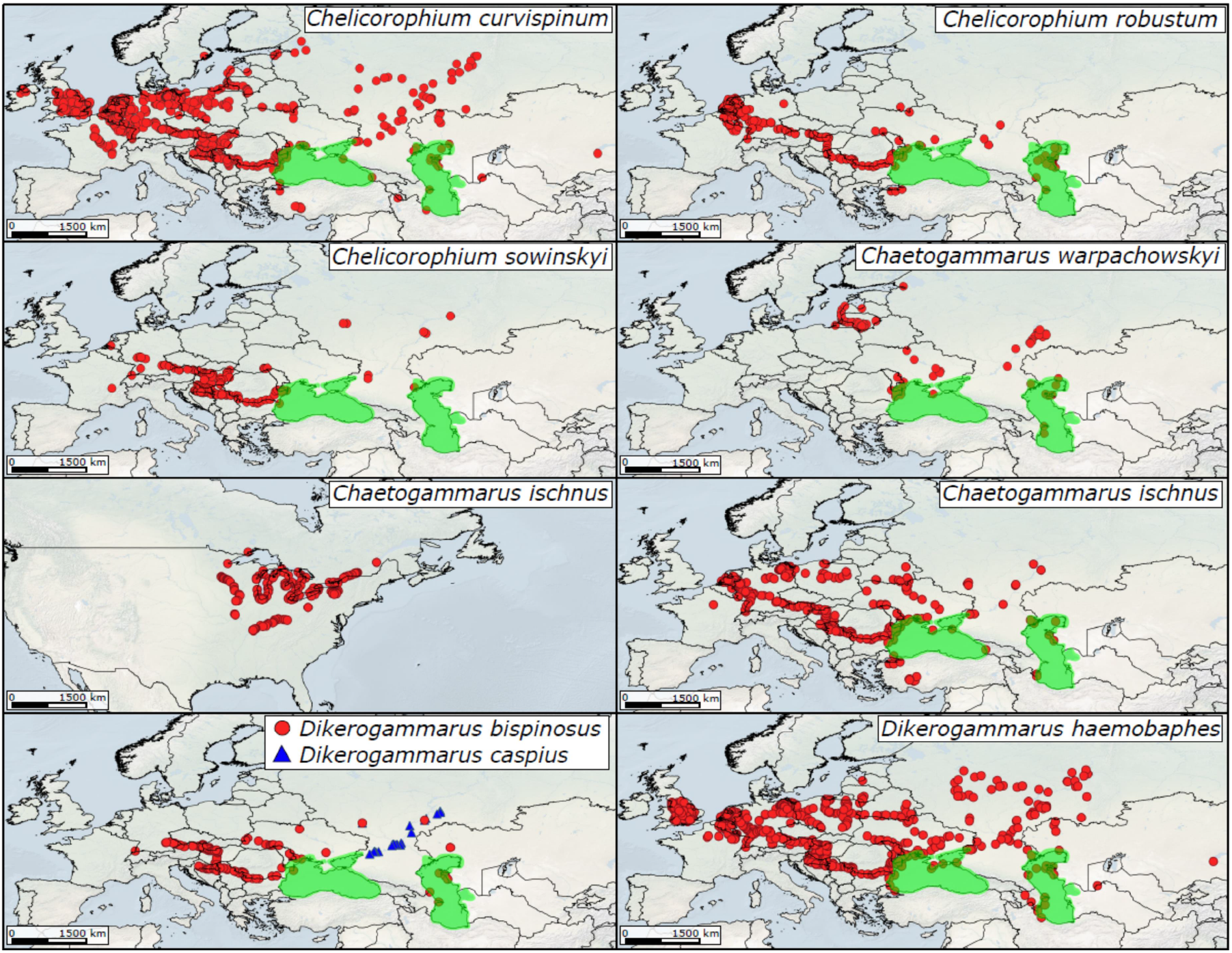
Distribution of widespread Ponto-Caspian amphipods in native and non-native areas (except *Dikerogammarus caspius* for which only non-native records are shown). Green shading denotes the native areal.

**Fig. 4.**
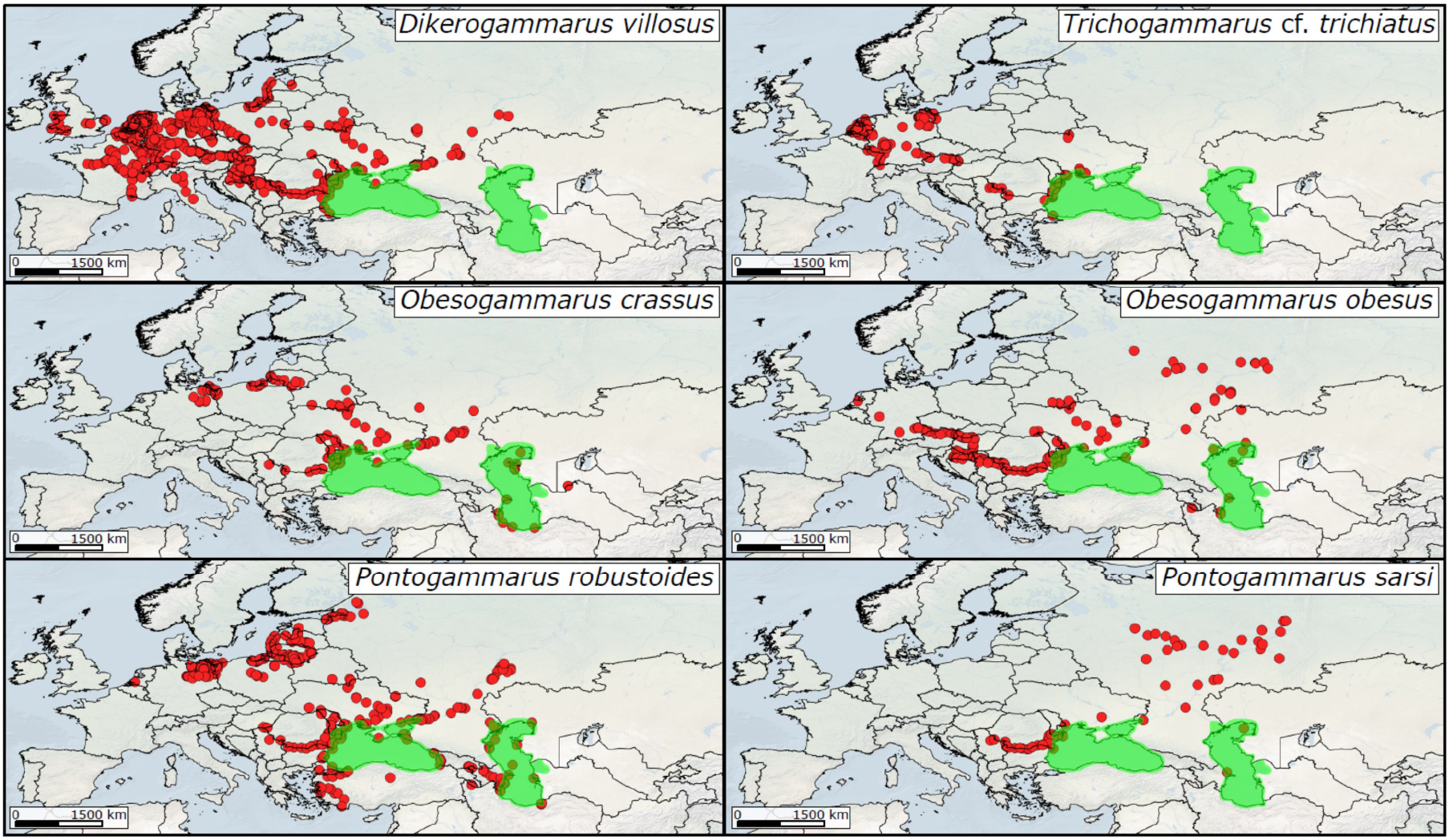
Distribution of widespread Ponto-Caspian amphipods in native and non-native areas. Green shading denotes the native areal.

**Fig. 5.**
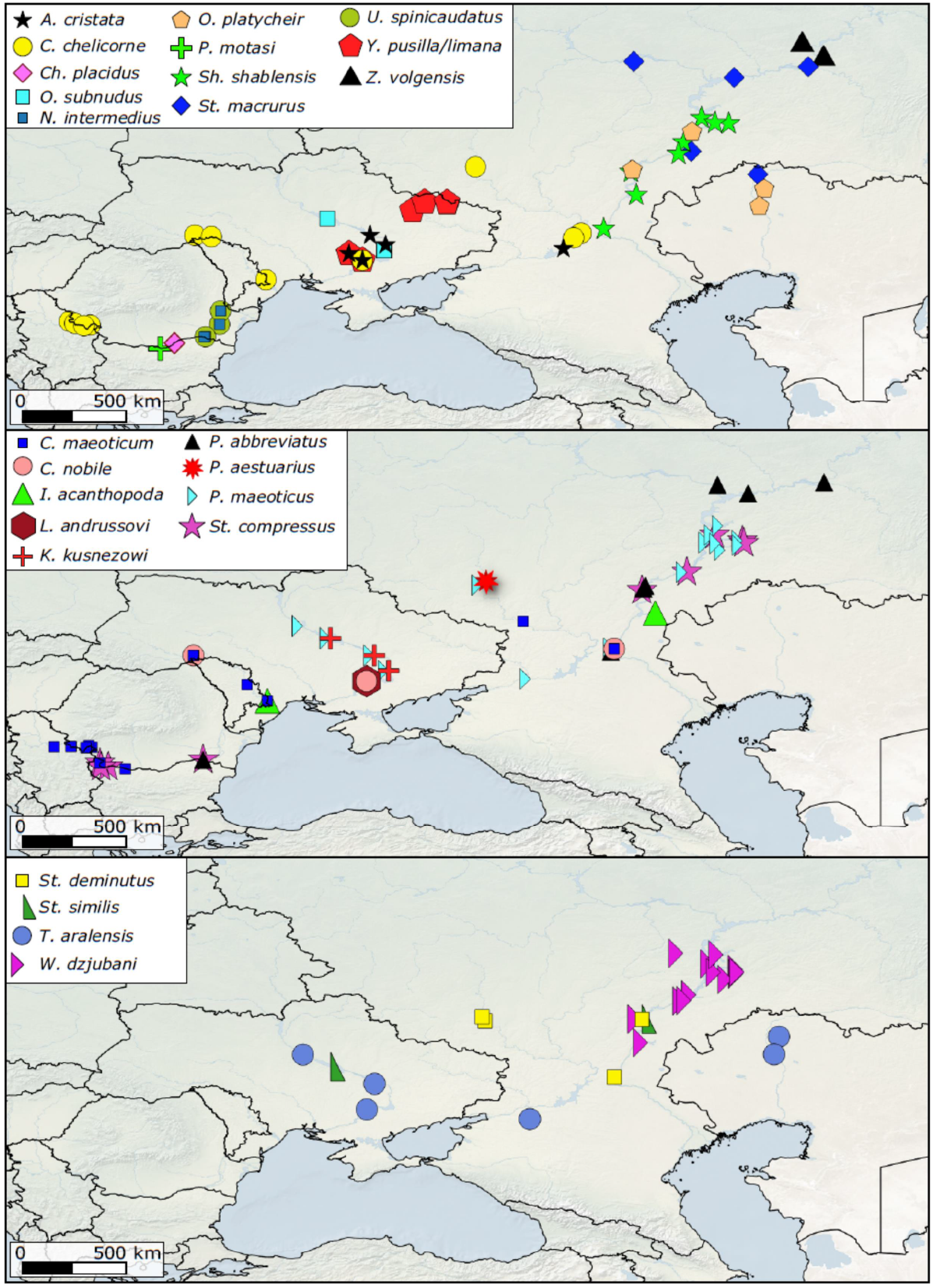
Distribution of Ponto-Caspian amphipods that have a less extended areal. All distribution points are non-native records.

Altogether, 34 countries spanning the Northern Hemisphere were found to harbor alien amphipods (Fig. 6A). The highest number of species was observed in countries adjacent to Ponto-Caspian water bodies such as Russia (30), Ukraine (24), Bulgaria (17), and Romania (16). A total of 25 freshwater ecoregions contained non-native species, the highest number being found in the ones closest to the Ponto-Caspian realm. These were the Volga-Ural (27 species), Dniester-Lower Danube (23), Dnieper-South Bug (22), and Don (20) (Fig. 6B).

**Fig. 6.**
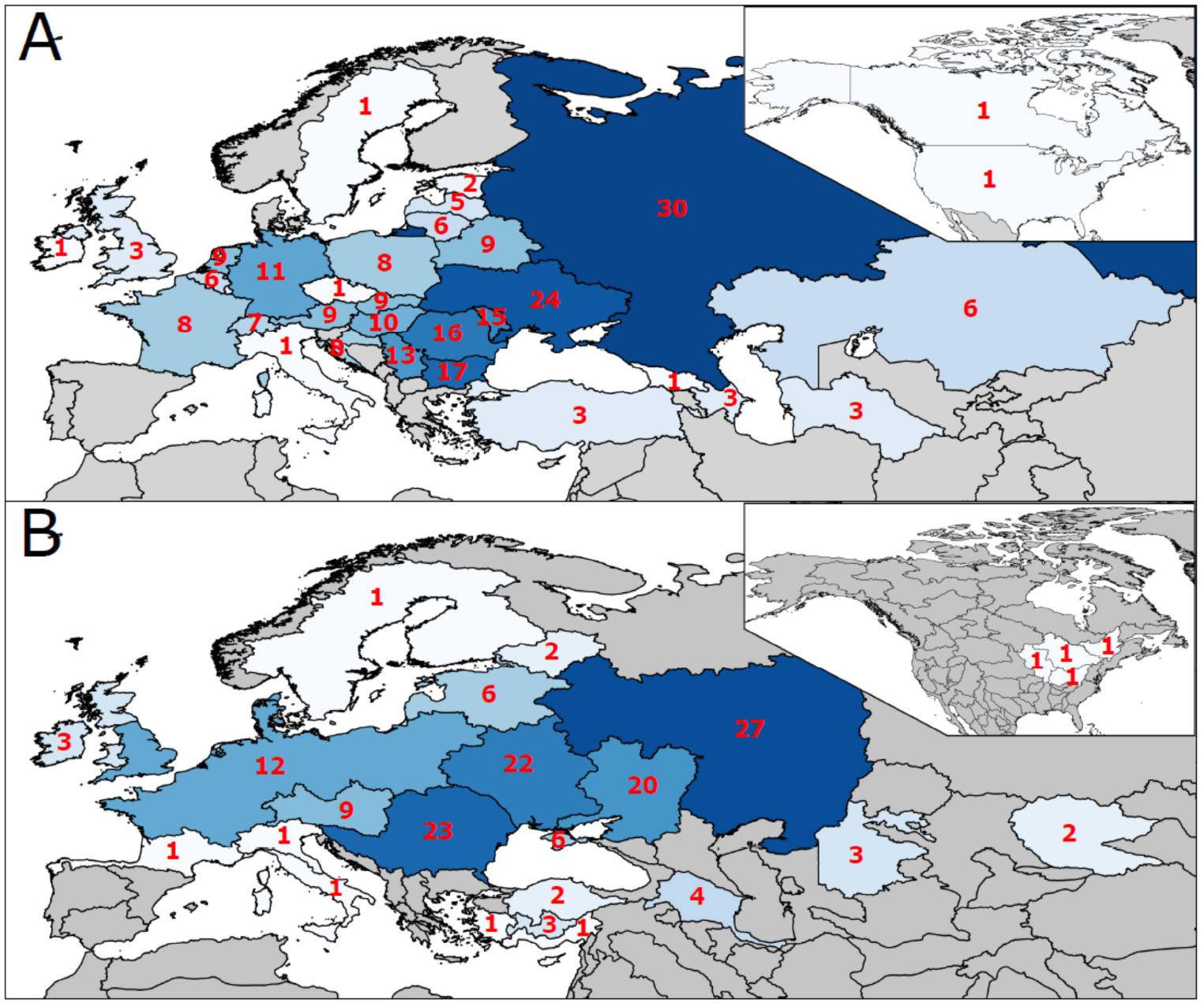
Number of alien Ponto-Caspian amphipod species per A) country and B) freshwater ecoregion.

Species richness analysis at a finer spatial scale revealed that the highest concentration of alien species is found throughout the lower courses of the largest rivers: Volga, Don, Dnieper and Danube. Generally, the number of alien species decreases gradually with distance from the native area (Fig. 7A). We also find that, in general, the sampling intensity is comparatively greater in the species-poor regions located further away from the Ponto-Caspian realm (Fig. 7B).

**Fig. 7.**
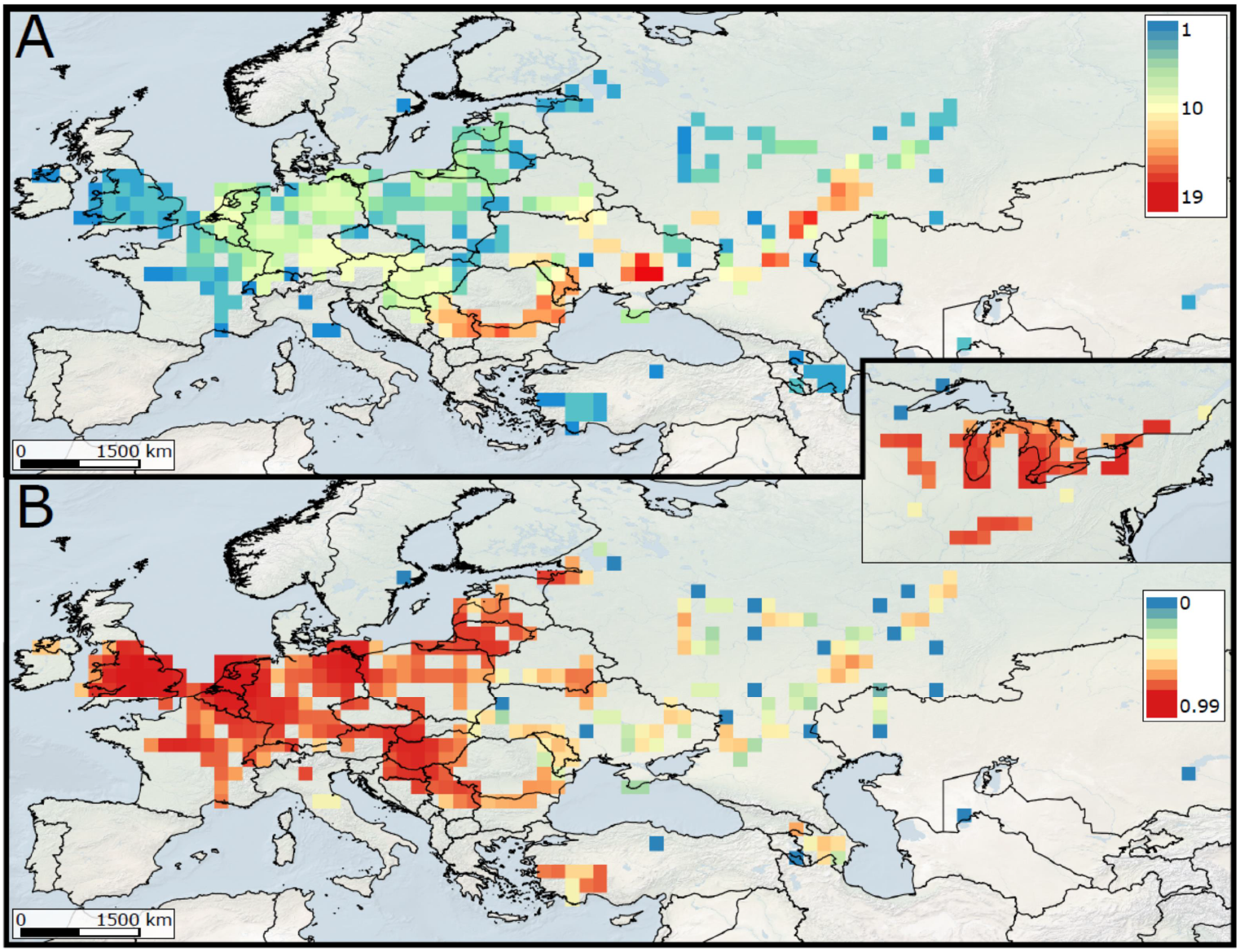
Species richness and sampling intensity of Ponto-Caspian amphipods in non-native areas. A) Distribution of species richness. Warmer colors indicate higher species number. B) Sampling redundancy. Warmer colors indicate higher sampling effort. Inset map represents the Great Lakes Region in North America.

Biogeographical clustering based on spatial species composition revealed three distinct faunistic regions (Fig. 8). Region 1 mainly encompasses the river drainages connected to the north-western part of the Black Sea (Rhine, Danube and Dniester) as well as a few areas of the upper-most reaches of the Volga basin. The most common species here is *C. curvispinum*. This region mainly corresponds to the Southern invasion corridor (Bij de Vaate et al. 2002). Region 2 encompasses the south and south-eastern edges of the Baltic Sea. The most common species is *P. robustoides*. It partially corresponds with the Central invasion corridor. Finally, region 3 comprises drainages that flow to the northern Black Sea, the Azov Sea and the Caspian Sea (Dnieper, Don and Volga). The most common species is *D. haemobaphes*. This region partially corresponds with the Northern invasion corridor.

**Fig. 8.**
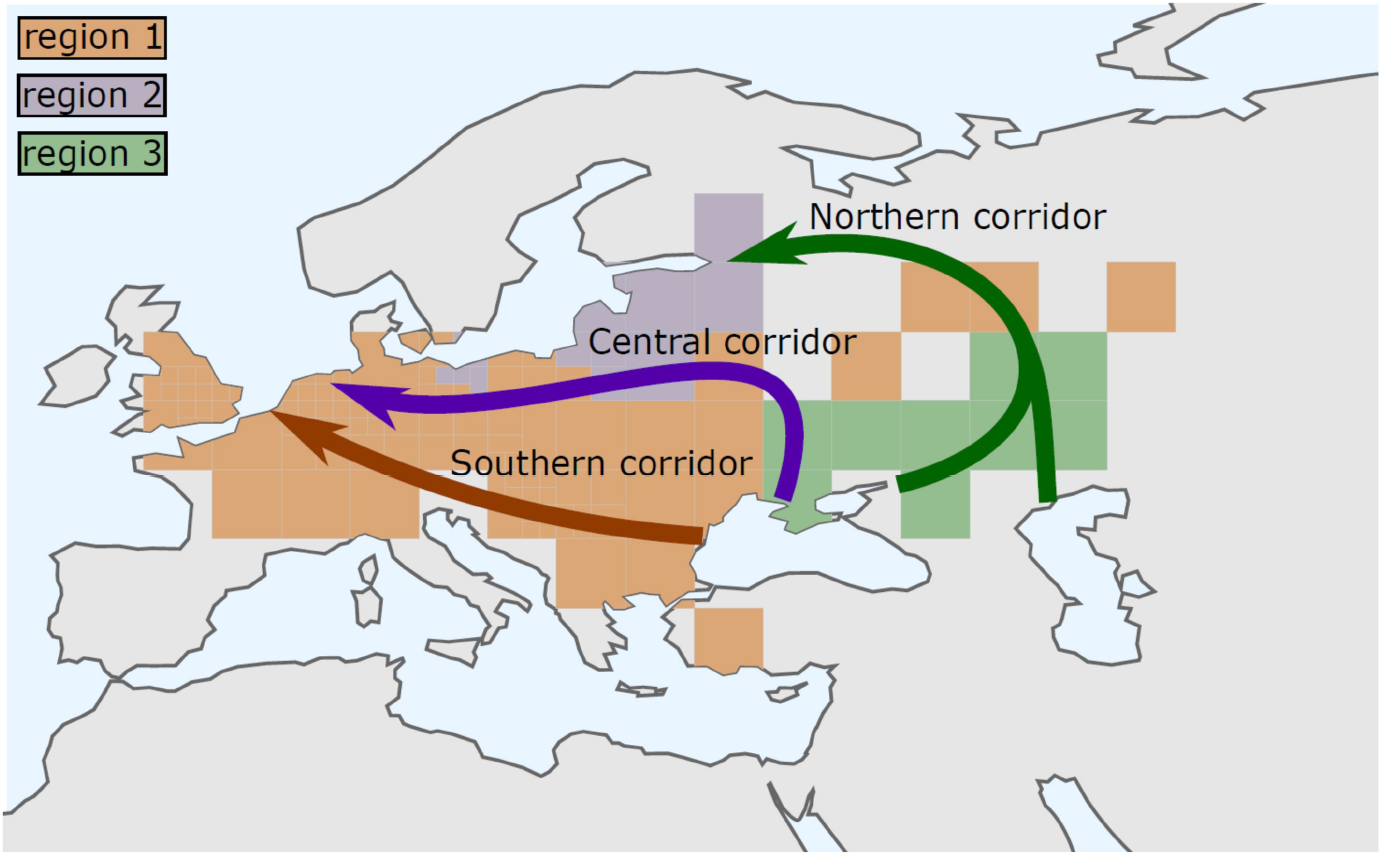
Biogeographical regions based on species composition. The colored arrows indicate the main invasion corridors through which Ponto-Caspian species have dispersed.

Additionally, we classified the geographical patterns of distribution into four categories (Table 1):

a. Type I - Species which are widespread in both Black Sea and Caspian basins, and their connected rivers (6 species).
b. Type II - Species mainly distributed in the Black Sea basin and connected rivers, with limited/no distribution in the Caspian basin (5 species).
c. Type III - Species mainly distributed in the Caspian basin with limited/no distribution in the Black Sea basin and connected rivers (4 species).
d. Type IV - Species with limited distribution in either or both basins (24 species). Therefore, we can summarize that 24 species (62%) have restricted distributions, six species (15%) are widespread, and nine species (23%) have intermediary range sizes.

## Discussion

Our study provides the first comprehensive overview of the diversity and distribution of APCAs at the global scale. We find that the non-native species are a representative subsample of the total Ponto-Caspian amphipod diversity in terms of taxonomy and ecomorphology. Our detailed, high-resolution distribution maps provide a first global perspective on the spatial extent of these species, emphasizing the role of anthropogenic corridors in shaping their non-native biogeography.

### Biodiversity

The total number of Ponto-Caspian amphipods that were found to occur outside the native region is quite high. Up to 39 species can be considered alien, representing about 40% of the entire endemic diversity. At higher taxonomic levels we find that 19 genera (out of 39, i.e. ∼49%) and five families (out of 10, i.e. 50%) contain alien species. It therefore appears that the non-native species represent approximately half of the entire endemic Ponto-Caspian amphipod taxonomic diversity. Furthermore, we also find that the alien species represent a good part of the total endemic phylogenetic diversity.

It appears that species rich genera have the highest number of alien species since we found a strong and positive relationship between the total number of species and alien species within genera. However, it is yet unclear whether this relationship reflects particular intrinsic generic traits that favor invasiveness, or is an artifact due to human activity. Perhaps these two factors are not mutually exclusive either.

From an ecomorphological perspective, our study reveals that the alien species belong to all of the four currently recognized Ponto-Caspian gammarid ecomorphs (Copilaș-Ciocianu and Sidorov 2021). Half of the species (49%) are diggers, and up to a quarter are crawlers (23%), while clingers and symbionts represent a small fraction (10% and 2%, respectively). These proportions reflect very well the situation in the native region, with the most common ecomorph being the digger (49%), followed by crawlers (26%), clingers (19%) and symbionts (6%) (Copilaș-Ciocianu and Sidorov 2021). Furthermore, we also did not find significant differences regarding the number of native and non-native species within ecomorphs. As such, we can deduce that alien species represent a significant subsample of the total ecomorphological diversity of endemic Ponto-Caspian amphipods. Further detailed morphometric analyses are necessary to paint a clearer picture regarding the morpho-functional differentiation (if any) among the native and alien species. Such insight can be useful in identifying the functional traits that favor invasiveness (Ordonez et al. 2010; Petruzzellis et al. 2021).

It remains to be seen whether the more numerous diggers and crawlers have certain advantages that promote invasiveness over clingers and symbionts. The first two are more associated with coarse substrates (fine sand to rocks) than the last two (Copilaș-Ciocianu and Sidorov 2021). An affinity for hard substrata has been proposed as an important factor favoring the dispersal of Ponto-Caspian peracarid crustaceans into heavily anthropized habitats (Borza et al. 2017). Indeed, numerous studies reported a preference of alien Ponto-Caspian amphipods for hard substrates, although relatively few species were studied thus far (Dermott et al. 1998; Hesselschwerdt et al. 2008; Jermacz et al. 2015; Borza et al. 2018; Poznańska-Kakareko et al. 2021). Clingers and symbionts seem to be associated with more specific substrates, which may limit dispersal potential. Therefore, our study also leans towards the hypothesis that preference for hard substrates is an important trait that characterizes non-native Ponto-Caspian crustaceans.

We consider that the introduction modes of many species are not yet certainly known. Most species have probably spread naturally, but this is only in conjunction with human activities, such as after being deliberately introduced, or after the construction of dams and canals (Grigorovich et al. 2002).

### Biogeography

Our high resolution maps capture for the first time the global spread of APCAs, providing a greater resolution and extent than any previous studies (Jazdzewski 1980; Bij de Vaate et al. 2002). We find that range sizes vary greatly among species, from highly localized to intercontinental. Nevertheless, despite the relatively large number of alien species, it appears that most of them (62%) have limited distributions. Only six species are widespread in both the Black Sea and Caspian basin, and nine are widely distributed only in one basin.

As expected, the highest alien species diversity is in the regions adjacent to the native areal, especially in the lower course of large rivers such as the Volga, Don, Dnieper and Danube. In Central Europe the highest diversity is observed in the Danube and Rhine rivers. Towards the outermost edges of the invaded areal the number of species is always low, these usually being the oldest invaders such as *C. curvispinum, C. ischnus* and *D. haemobaphes*.

Lower sampling intensity in areas close to the native areal suggests that more intensive monitoring is required here, as well as in the native region. This is especially important since many introductions took place among these regions, and most of the species diversity is found here (Grigorovich et al. 2002). Despite a few more recent studies (Uzunova 1999; Petrescu 2009; Zinchenko and Kurina 2011; Konopacka et al. 2014; Copilaș-Ciocianu et al. 2020; Kurina 2020; Son et al. 2020), there is a significant lack of new data in many of these areas. It is very likely that some of the introduced species are extinct, or their distributions could have changed significantly.

Biogeographical clustering of species composition revealed three faunistic provinces that correspond remarkably with the Northern, Central and Southern invasion corridors used by the Ponto-Caspian fauna (Bij de Vaate et al. 2002). These corridors consist of artificially connected large rivers and have limited to no connectivity among each other. This overlap between corridors and faunistic provinces highlights, on one hand, that human mediated dispersal plays a crucial role in the spreading of Ponto-Caspian fauna outside its native area. On the other hand, this overlap also emphasizes that the distinct faunal composition between region 1 (corresponding to the Southern corridor with species originating from the Black Sea) and region 3 (corresponding to the Northern corridor with species originating from the Azov and Caspian Seas) is connected to the biogeography of the area, since the Black and Caspian Seas have distinct biotas (Mordukhai-Boltovskoi 1964, 1979; Copilaș-Ciocianu and Sidorov 2021). Moreover, our results show that the Azov drainages have a greater similarity in alien species composition to the Caspian Sea rather than to the nearby Black Sea. This similarity is sometimes reflected in phylogeographic studies which identified a closer genetic connection of the Azov with the Caspian rather than the Black Sea (Audzijonyte et al. 2006; Nahavandi et al. 2013), most likely reflecting the region’s geological past (Krijgsman et al. 2019; Palcu et al. 2021). As such, it appears that the regional species pools in the native areal coupled with a strong anthropogenic influence play a critical role in shaping the non-native distributions of Ponto-Caspian amphipods.

### Prospects for further dispersal

Europe and temperate Asia are the most affected regions by alien crustacean species, and their number is predicted to significantly rise by 2050 (Seebens et al. 2021). Borza et al. (2017) concluded that the possibility for additional Ponto-Caspian peracarid crustaceans of becoming invasive in the future is low. Nevertheless, given that the current number of non-native species is already quite high, further dispersal into new regions is unavoidable. This is exemplified by the regular reporting of new APCAs records (e.g. Csabai et al. 2020; Son et al. 2020; Lipinskaya et al. 2021). As such, additional monitoring is needed, coupled with the implementation of management measures aimed at minimizing further dispersal (Vander Zanden and Olden 2008).

It remains to be seen how much further APCAs will spread in the future. Although local range expansions are constantly being reported, most of them occur in Europe. Eastward dispersal into large Siberian rivers such as Ob and Yenisei is likely, as this pattern has been recently observed in pond mussels and was connected to fish stocking (Kondakov et al. 2020). The species with the most eastward occurrence are *C. curvispinum* and *D. haemobaphes*, being present in the delta of the Ile River and some parts of Lake Balkhash (Petr 1992; Khassengaziyeva and Mamilov 2020). Considering that these two species are also the most widespread of all Ponto-Caspian amphipods, they are the prime candidates for being detected in the large Siberian rivers in the future. However, the possibility of jump dispersal due to shipping activity make future predictions less straightforward given the apparent haphazard nature of these events (Copilaș-Ciocianu et al. 2020). Nevertheless, climatically suitable areas that may support such populations can be identified on the basis of modeling.

Our high resolution distribution data will be invaluable for modeling future distributions and predicting future invasion pathways of APCAs (Mainali et al. 2015; but see Liu et al. 2020). This is especially critical under future climate warming scenarios (Zhang et al. 2020). Predicting which invasive species will spread further and which native species will go extinct or decline is of major importance for implementing adequate conservation measures (Porfirio et al. 2014). Another important aspect of our data is that it will also allow testing whether APCAs experience climatic niche shifts and/or expansions in the non-native areal relative to their native areal (Broennimann et al. 2007; Guisan et al. 2014). This will be critical for assessing long-term dynamics, persistence, and adaptation of APCAs in non-native habitats.

## Acknowledgements

This study was financed by the Research Council of Lithuania (contract no. S-MIP-20-26). We are grateful to Nadezhda Berezina, Péter Borza, and Sonia Uzunova for providing important literature.

## Declarations

### Availability of data and material (data transparency)

Data is available at Figshare.

### Authors’ contributions

DCC, ESC and DS—conceptualization, DCC, ESC and DS—literature screening, DCC—data collection and formal analysis, DCC—led the writing; all authors approved the final version of the manuscript.

